# Genome-Wide CRISPRi Screening Identifies XPO5 as a Regulator of B Cell Mutation and Fitness

**DOI:** 10.64898/2026.04.22.720139

**Authors:** Tylar G. Kirch, Zachary D. Miller, Jacob S. Dearborn, William G. Dowell, Sylvester Languon, Hana Paculova, Jade H. Cleary, Seth E. Frietze, Kalev Freeman, Devdoot Majumdar

## Abstract

The B cell receptor (BCR) is the defining factor of B lymphocyte identity and function, allowing for a robust adaptive immune response through antigen recognition. Strict regulation of BCR surface density dictates proper B cell signaling, immune regulation, and the prevention of malignancy, yet the factors regulating this density remain undefined. Here, we performed a genome-wide CRISPR interference (CRISPRi) screen in Ramos B cells, which undergo constitutive somatic hypermutation (SHM) and identified Exportin-5 (XPO5) as a central regulator of BCR surface expression. XPO5 depleted cells exhibited an accelerated loss of surface BCR with no change in transcript levels, suggesting a potential post-transcriptional regulatory mechanism. Further analysis revealed XPO5 depletion led to an accumulation of non-functional BCR light chain sequences driven by an increase in AID signature mutations, implicating XPO5 in balancing mutagenesis and repair during somatic hypermutation (SHM). Transcriptomic and small RNA sequencing revealed a global reduction in miRNA levels and enrichment of target gene sets indicative of cell cycle arrest and increased DNA damage response. These data suggest that XPO5 plays a multi-faceted regulatory role in B cells via a miRNA-mediated control, supporting both proliferation and regulating DNA repair thresholds to maintain B cell receptor expression and functionality.

**Significance:** Precise regulation of B cell receptor (BCR) density is essential for immune function and preventing malignancy. Through a genome-wide CRISPR interference (CRISPRi) screen, we identified Exportin-5 (XPO5) as a critical regulator of BCR surface expression. We show that XPO5 is essential to maintain the miRNA landscape that supports DNA repair during somatic hypermutation. Loss of XPO5 destabilizes this mutational balance, driving the accumulation of non-functional BCR sequences. This study uncovers a novel connection between miRNA nuclear export and the preservation of B cell identity and genomic fidelity, highlighting the multi-faceted regulatory role of XPO5.

## Introduction

B lymphocytes are central to the adaptive immune response, primarily acting through the production of antibodies. The membrane bound form of an antibody, known as the B cell receptor (BCR), dictates nearly all core functions of B cell biology and acts as a central dependency in many B cell malignancies. The BCR facilitates antigen recognition and transduces signals crucial to B cell survival, proliferation, and differentiation.^1–3^ In the absence of antigen, B cells rely on low level ‘tonic’ signals to maintain general cell survival and homeostasis.^4–6^ The BCR serves as the master regulator of B cell identity, survival, and ultimately a robust adaptive immune response.

The number of BCR molecules at the cell surface is highly regulated throughout B cell stimulation and development. Elevated BCR surface density amplifies antigen-dependent signaling strength, which drives germinal center (GC) selection dictating downstream fate decisions such as the development into memory or plasma B cells.^7–9^ Additionally, high BCR surface density can enhance antigen-independent ‘tonic’ signaling, further increasing the likelihood of B cell survival and positive selection.^10^

Disruption or the dysregulation of BCR surface density is a hallmark of malignancy and autoimmunity. Chronic, dysregulated BCR signaling fuels the pathogenesis of multiple B cell cancers, including chronic lymphocytic leukemia (CLL)^11,12^, mantle cell lymphoma (MCL)^13,14^, Burkitt lymphoma (BL)^15,16^, and subtypes of diffuse large B cell lymphoma (DLBCL).^17,18^ In these contexts, high BCR surface density hyperactivates multiple oncogenic signaling pathways such as NF-κB, MAPK, and mTOR promoting uncontrolled proliferation and survival.^18–21^ Conversely, insufficient BCR surface density impairs negative selection, allowing autoreactive B cells to escape tolerance checkpoints to promote autoimmunity.^22,23^ Thus, the precise control of BCR surface levels is indispensable for proper signaling thresholds and effective humoral immunity.

Although transcriptional programs have traditionally been the primary focus in studies of immune regulation, growing evidence highlights the necessity of highly regulated post-transcriptional mechanisms. RNA-binding proteins (RBPs) and splicing factors regulate B cell development and function by controlling mRNA processing, stability, and translation.^24–26^ For instance, PTBP1 drives the selection of high-affinity B cell clones by regulating splicing events that control c-MYC-dependent proliferation and survival.^27^ Within the HNRNP family, HNRNPLL and HNRNPF regulate splicing during plasma cell differentiation and germinal center responses, respectively.^28,29^ Even the expression of the BCR itself is post-transcriptionally regulated by ZFP318, mediating an alternative splicing event that allows naïve B cells to co-express IgM and IgD BCR isotypes.^30^ Despite these insights, the specific post-transcriptional mechanisms that maintain precise control of BCR surface density remain incompletely defined.^31–39^

Here, we systematically identify regulators of BCR surface expression using a genome-wide CRISPR interference (CRISPRi) screen in B cells. This approach identified Exportin-5 (XPO5), a component of nuclear RNA export and miRNA biogenesis machinery^40–42^, as a factor required for sustained BCR surface expression. We demonstrate that XPO5 is a multifunctional regulator in B cells responsible for preserving both BCR expression and cellular homeostasis. The depletion of XPO5 results in progressive loss of surface BCR driven by a failure in DNA repair mechanisms and proliferation defects.

## Results

### Genome-wide CRISPRi screen identifies XPO5 as a regulator of surface BCR expression in Ramos B cells

We performed a genome-wide pooled CRISPR interference (CRISPRi) screen in Ramos cells, a Burkitt lymphoma-derived line that stably expresses the IgM isotype BCR and is widely used as a model of B cell malignancy, to identify regulators of BCR expression. Ramos clones stably expressing dCas9-KRAB were transduced with the human CRISPRi-V2 sgRNA library^43^, containing 209,070 unique single guide RNA (sgRNA) sequences targeting 18,905 genes, including additional non-targeting controls at a low MOI (∼0.1) to ensure single-sgRNA integration. Transduced cells were selected by puromycin resistance and after 14 days in culture BCR^pos^ and BCR^neg^ cell populations were isolated by flow cytometry, alongside an unsorted control (presort). Cell numbers were maintained at high coverage (≥100x) to ensure uniform sgRNA representation. Genomic DNA from each group was sequenced to quantify sgRNA representation and identify genes whose perturbation altered surface BCR expression (Fig 1A).

**Figure 1.**
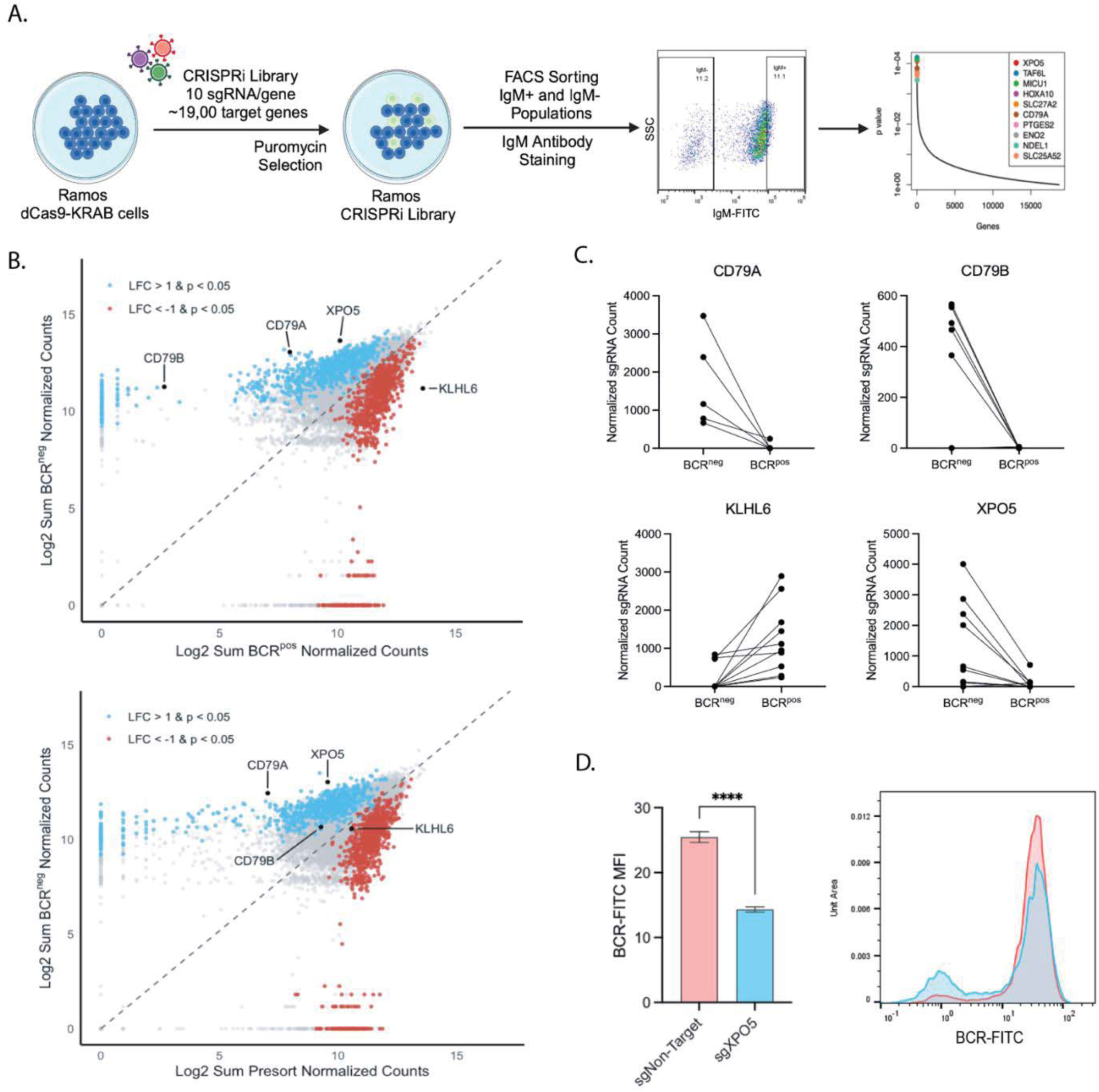
Genome-wide CRISPRi screen identifies XPO5 as a regulator of surface BCR expression in Ramos B cells. (A) Schematic workflow of genome-wide CRISPRi screen. Ramos B cells stably expressing dCas9-KRAB were transduced with a genome-wide sgRNA library (10 sgRNA per gene) and subjected to puromycin selection. Cells were stained for surface IgM and sorted via fluorescence activated cell sorting (FACS) into BCR negative (BCR^neg^) and BCR positive (BCR^pos^) populations. sgRNA abundance in sorted populations was determined by next-generation sequencing to identify gene hits ranked by p-value. (B) Scatter plots displaying the comparison of normalized sgRNA counts (in log2) between BCR^neg^ and BCR^pos^ sorted populations (top) and between BCR^neg^ and presort populations (bottom). Genes significantly enriched in the BCR^neg^ population are colored blue (log2 fold change >1; p < 0.05), while significantly depleted genes are colored red (log2 fold change <-1; p < 0.05). Known BCR positive (*CD79A*, *CD79B*) and negative (*KLHL6*) regulators and identified hit *XPO5* are highlighted. Significance thresholds and p-values were determined using the MAGeCK RRA (Robust Rank Aggregation) algorithm. (C) Ladder plots showing normalized counts for individual sgRNAs targeting highlighted genes (*CD79A*, *CD79B*, *KLHL6*, and *XPO5*) between BCR^neg^ and BCR^pos^ sorted populations. Each line compares individual sgRNA enrichment between populations. (D) Validation of XPO5 knockdown phenotype. Ramos dCas9-KRAB cells were transduced with individual sgRNA targeting XPO5 (sgXPO5) or a non-targeting control (sgNon-Target) and surface BCR was determined by flow cytometry. (Left) Quantification of BCR-FITC mean fluorescence intensity (MFI). Data presented as mean ± S.D. of n=6 biological replicates. ****p < 0.0001 determined by t-test. (Right) Corresponding flow cytometry histogram showing shift in surface BCR expression between sgNon-Target (red) and sgXPO5 (blue) cells.

Enrichment analysis of sgRNA sequences in the BCR^neg^ population (where positive regulators of BCR expression are expected) compared to both presort and BCR^pos^ populations identified several bona fide regulators of surface BCR. For instance, the well-characterized adapters of BCR expression CD79A and CD79B are well known to be required for surface expression of BCR and serve as signal transducers of BCR signaling; fittingly, both proteins feature prominently in the top 0.5% of hits (Fig. 1B-C).^44,45^ Conversely, sgRNAs targeting KLHL6, a known negative regulator of CD79, were enriched in the BCR^pos^ population (Fig. 1B-C).^46,47^ Unexpectedly, a prominent exporter of pre-miRNAs, XPO5 featured as the top ranked candidate enriched in the BCR^neg^ population, suggesting a role as a positive regulator of BCR expression (Fig. 1B-C).

Next, we validated these putative regulators independently with individual sgRNAs. The top enriched sgRNA sequence targeting XPO5 resulted in a reduction in BCR surface expression (Fig. 1D *left*). Curiously, this reduction was primarily observed as an emergence of a distinct BCR negative population rather than a shift in surface density across the population (Fig. 1D *right*). While general miRNA processing is known to be essential for B cells function and development, the specific contribution of XPO5 in B cell biology and its role in maintaining BCR surface expression remains undefined.^48,49^

### XPO5 knockdown specifically reduces surface BCR expression

To investigate the mechanism driving the observed increase in the BCR^neg^ population, we first confirmed XPO5 knockdown efficiency by RNA sequencing. We observed a 76.7% reduction in XPO5 expression with a corresponding log2 fold change of -2.0 (Fig. 2A-B). However, despite the decrease of BCR surface expression, we detected no reduction in transcript levels of BCR heavy chain (*IGHM*), light chain (*IGLC*), or BCR signaling adaptors required for surface expression *CD79A* and *CD79B* (Fig. 2A-B). This decoupling of BCR mRNA from surface protein expression suggests the regulatory role of XPO5 is unlikely to be transcriptional and might be the result of a post-transcriptional mechanism.

**Figure 2.**
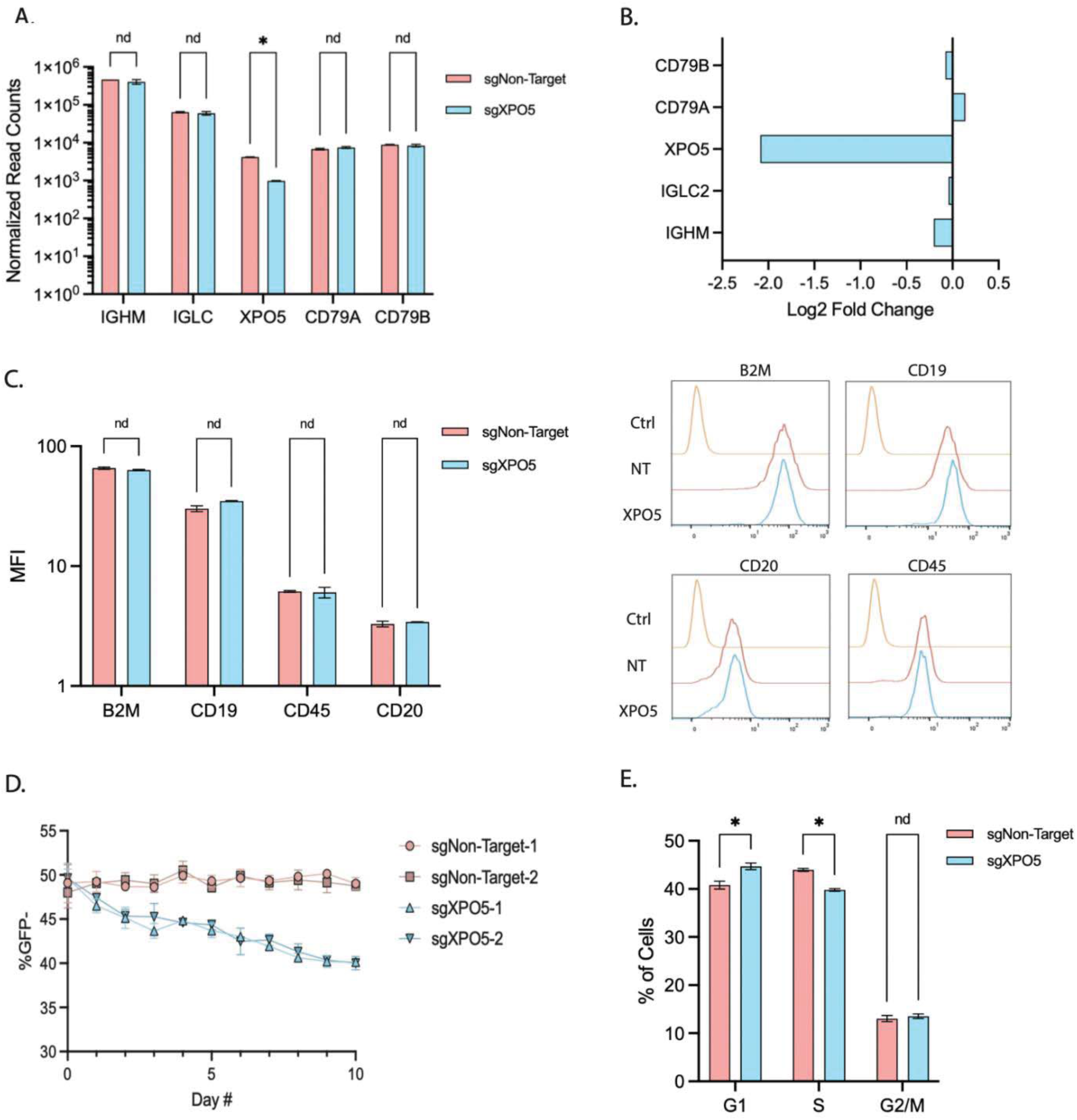
XPO5 knockdown specifically reduces surface IgM expression. (A) Normalized RNA-seq read counts for BCR components (IGHM, IGLC, XPO5, CD79A, CD79B) and XPO5 in dCas9-KRAB expressing Ramos cells transduced with sgRNA targeting XPO5 (blue) or non-target control (Red). Data presented as mean ± S.D. of n=2 biological duplicates. nd, no significant difference; *p < 0.05 determined by t-test (B) Corresponding log2 fold changes of transcripts shown in (A). (C) Flow cytometry analysis of surface expression specificity. Mean Fluorescence Intensity (MFI) of B2M. CD19, CD45, and CD20 (mean ± S.D., n=3) (Left). Representative histograms showing unstained control (orange), sgNon-Target (red) and sgXPO5 (blue) (right). (D) Growth competition assay. GFP negative XPO5 knockdown cells were mixed with GFP expressing non-targeting control cells and monitored over 10 days. Data presented as mean ± S.D. of n=2 biological replicates. (E) Cell cycle analysis showing distribution of cells in G1, S, and G2/M phases. Data presented as mean ± S.D. of n=3 biological replicates. nd, no significant difference; *p < 0.05 determined by t-test.

Next, we determined if XPO5 acts as a specific regulator of BCR or a general modulator of membrane protein expression. In order to differentiate between these possibilities, we examined other commonly expressed B cell surface proteins. Monitoring surface expression levels of MHC Class I component β2-Microglobulin, and common surface markers CD19, CD45, and CD20 revealed no significant changes as a function of XPO5 expression (Fig. 2C). This data suggests XPO5 is not acting as a regulator of global membrane protein expression.

Given XPO5 elicits either oncogenic or tumor-suppressive roles in numerous cancer types^50–54^, we next investigated its role in Ramos B cell proliferation and fitness. We performed a competition assay, as previously described elsewhere^44^, co-culturing XPO5 depleted and GFP expressing non-targeting control cells at a 1:1 ratio. Competing cells were monitored for 10 days to permit a large dynamic range of fitness defects. At assay endpoint, the absence of XPO5 resulted in a marked decrease in cell numbers relative to controls, suggesting a necessity for XPO5 in maintaining optimal proliferation rates (Fig. 2D). To better understand the defect in cell fitness, we performed cell cycle analysis. XPO5 depletion led to an increased proportion of cells in the G1 phase and a corresponding decrease in S phase (Fig. 2E). These findings suggest XPO5 is required for efficient cell cycle progression, specifically at the G1/S phase transition.

### XPO5 knockdown accelerates the loss of BCR expression

Noticing a prominent BCR^neg^ population in Fig 2A, we then focused on the ontogeny of this population. One source of the BCR^neg^ population is due to the propensity of Ramos cells to undergo continuous somatic hypermutation (SHM), a steady background of BCR^neg^ cells is common in long-term experiments.^55–57^ In order to properly account for the role of background levels of SHM in the Ramos B cell line, we next conducted experiments to measure levels of the BCR^neg^ population. Surprisingly, we found that XPO5 knockdown significantly accelerated the increase in BCR^neg^ cell percentages to ∼15% over controls after 20 days (Fig. 3A-B). This increase in the BCR^neg^ population suggested the possibility that XPO5 may regulate the rate of SHM or downstream repair mechanisms.

**Figure 3.**
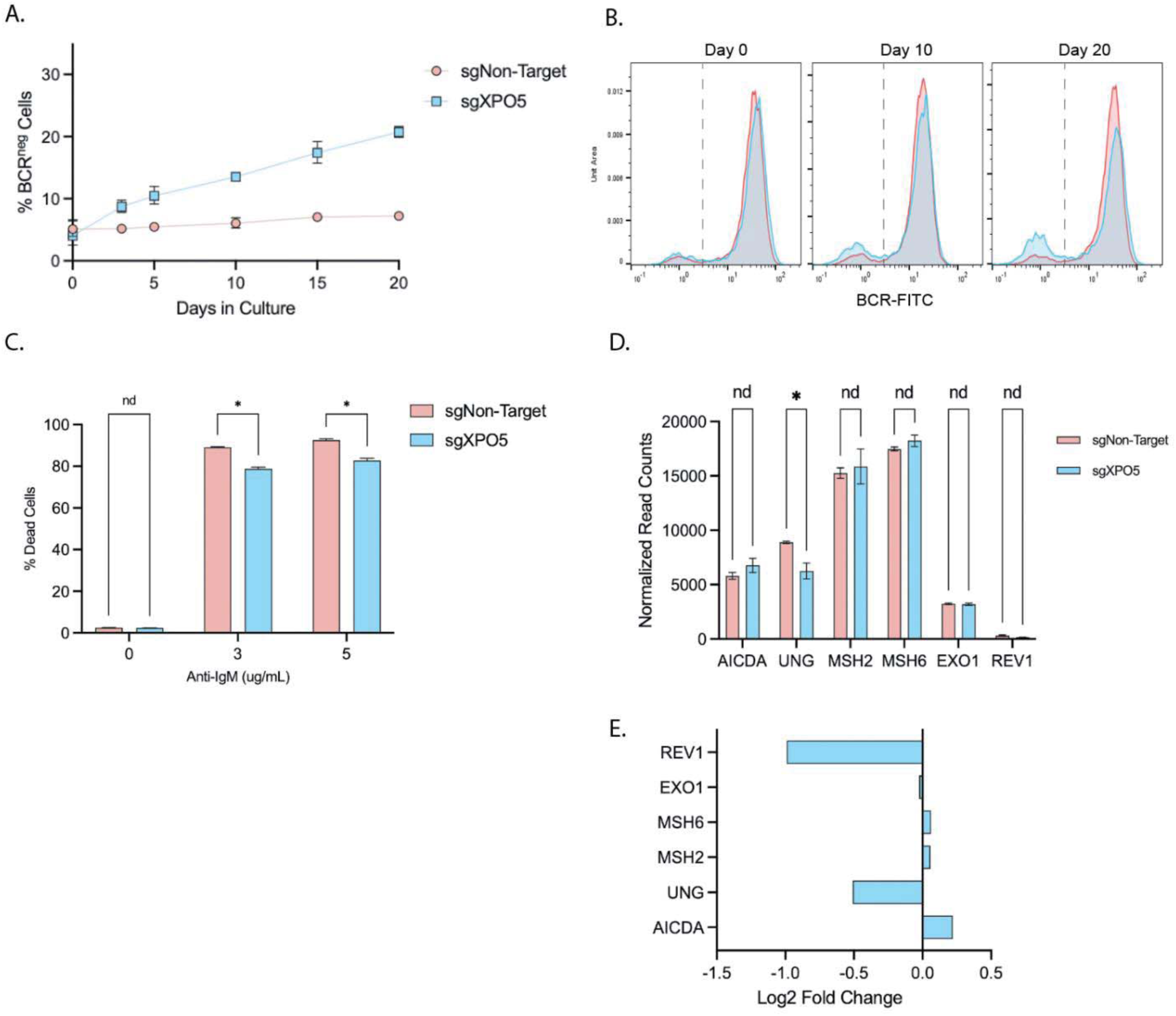
XPO5 knockdown accelerates the loss of BCR expression. (A) Time course analysis of surface BCR loss. The percentage of BCR^neg^ cells were monitored by flow cytometry over 20 days. sgXPO5 cells (blue) lose surface BCR at a higher rate than non-target controls (red). Data presented as mean ± S.D. of n=3 biological replicates. (B) Representative flow cytometry histograms of surface IgM expression at Day 0, 10, and 20. The dashed vertical line indicates the gate used to define the BCR^neg^ population quantified in (A). sgXPO5 populations (blue) show a progressive accumulation in the negative gate compared to non-target controls (red). (C) Anti-BCR (IgM) induced apoptosis assay. Percent cell death in sgNon-Target (red) and sgXPO5 (blue) cells following 24-hour treatment with increasing concentrations of anti-IgM (0, 3, 5 μg/mL). Data presented as mean ± S.D. of n=3 biological replicates. nd, no significant difference; *p < 0.05 determined by t-test. (D) Normalized read counts for somatic hypermutation (SHM) and DNA repair factors (*AICDA*, *UNG*, *MSH2*, *MSH6*, *EXO1*, and *REV1*) derived from RNA-seq. Data presented as mean ± S.D. of n=2 biological replicates. nd, no significant difference; *p < 0.05 determined by t-test. (E) Corresponding log2 fold changes of the transcripts shown in (D).

We hypothesized that as the population of BCR^neg^ cells increased in XPO5 depleted cells, the overall population would be less susceptible to BCR mediated apoptosis. To test this, we utilized the anti-IgM stimulation assay, leveraging the known mechanism that BCR cross linking induces apoptosis in Ramos cells.^58,59^ After 24 hours of BCR stimulation, XPO5 depleted cells exhibited a ∼10% increase in resistance compared to controls, confirming the BCR^neg^ population has a deficiency in BCR signaling capability (Fig. 3C).

To examine possible effects of XPO5 depletion in increasing SHM, we examined transcriptional levels of related proteins. Transcriptional analysis revealed that depletion of XPO5 resulted in a significant 29.7% decrease in *UNG* expression, which initiates base excision repair by excising uracil and a 49.6% decrease in *REV1*, a DNA polymerase involved in translesion synthesis.^60,61^ We also observed a 16.5% increase in *AICDA* expression, although not significant, this trend may be notable given the tight regulation of AID in B cells (Fig. 3D).^62–64^ These changes in expression corresponded to log2 fold changes of approximately -0.5, -1.0, and 0.22 for *UNG*, *REV1*, and *AICDA* respectively, while no significant transcriptional changes occurred in other SHM associated repair pathways (Fig. 3E). Together, these data are consistent with altered SHM-repair dynamics following XPO5 depletion.

### XPO5 knockdown increases mutation frequencies in BCR light chain

Based on the observed transcriptional changes in DNA repair genes, we hypothesized that the accelerated accumulation of BCR^neg^ cells might be driven by an increase in mutations at the BCR locus. Using a TRUST4 pipeline^65^, RNAseq reads were aligned to the dominant BCR sequence for both heavy and light chain to analyze mutation frequencies. We primarily focused on the variable segments of the BCR as this is the primary target site for AID as well as disruption to this region being the primary cause of BCR^neg^ Ramos cells. Analysis of BCR sequence functionality revealed a significant increase in the ratio of non-functional to functional light chain BCR sequences, characterized by premature stop codons (Fig. 4A). The increase in non-functional light chain sequences indicates a potential loss of BCR functional integrity to limit surface expression.

**Figure 4.**
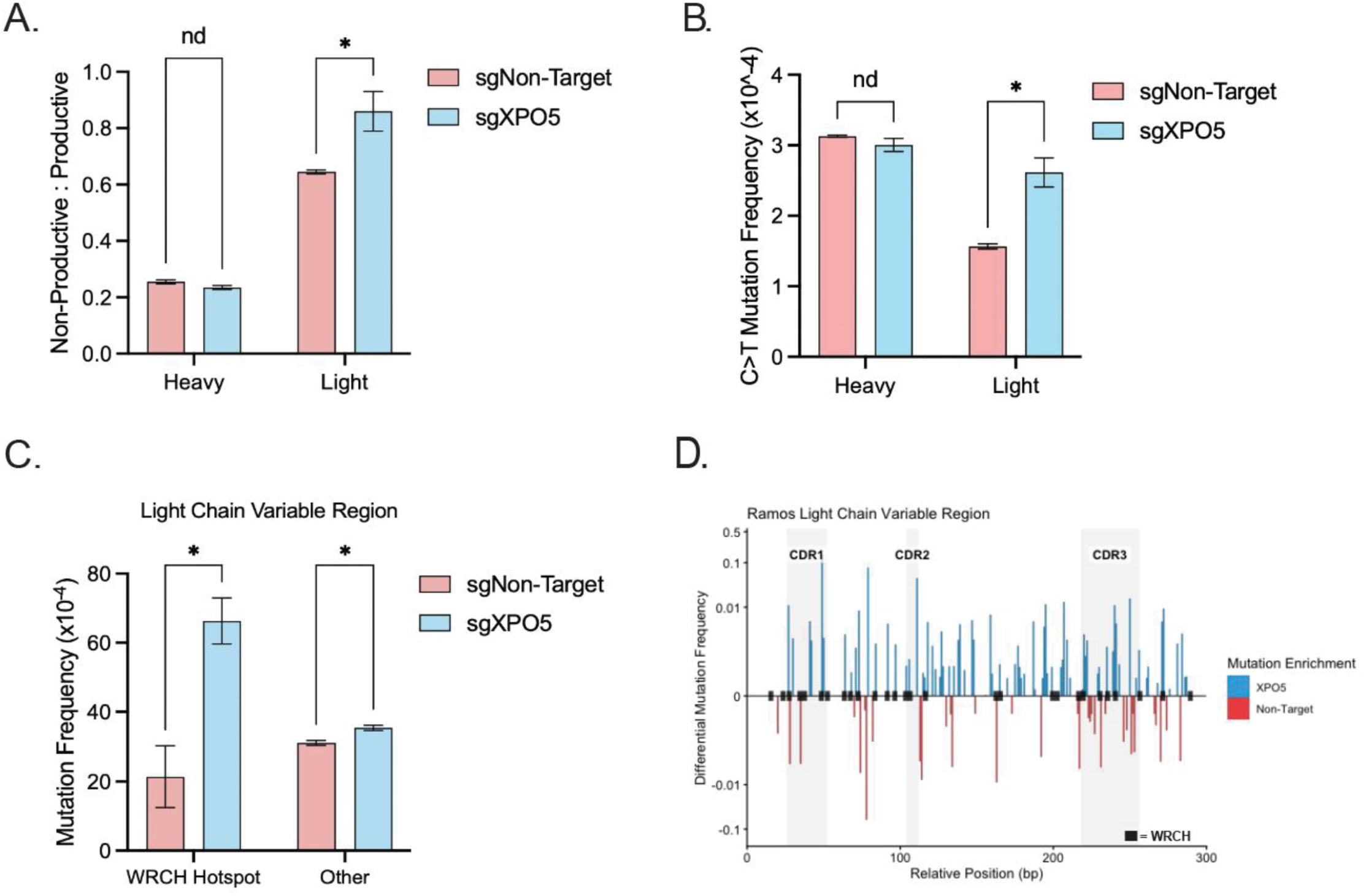
XPO5 knockdown increases mutation frequencies in BCR light chain. (A) Ratio of non-productive to productive BCR sequences heavy and light chains. sgXPO5 (blue) shows increased non-functional light chain sequences compared to sgNon-Target (red). (B) Frequency of C-to-T transition mutations within the heavy and light chain variable regions. The specific increase in C>T transitions in the light chain of sgXPO5 cells indicates a failure to repair AID-generated uracil lesions. (C) Mutation frequencies in the light chain stratified by canonical AID targeting motifs (WRCH hotspots) versus non-hotspot sequences (Other). Increased mutagenesis in sgXPO5 cells is preferentially targeted to AID hotspots (D) Mutation analysis of the light chain VJ regions. Differential mutation frequencies at each nucleotide position for sgXPO5 (blue, top) and sgNon-Target (red, bottom). WRCH motifs are marked with black squares along axis. All data presented as mean ± S.D. of n=2 biological replicates. nd, no significant difference; *p < 0.05 determined by t-test.

To determine if this loss of function stemmed from a specific failure in DNA repair pathways, we analyzed the mutational spectrum of these sequences. AID initiates SHM by deaminating cytosine creating uracil lesions, while these uracils are excised by UNG to allow high fidelity repair or error prone processing.^57,61,62^ Strikingly, XPO5 depleted cells exhibited a significant enrichment of C to T transition mutations within the light chain (Fig. 4B). Since C to T mutations result from the direct replication of unexcised uracils, this specific mutational signature indicates a failure in UNG mediated base excision, which is consistent with the transcriptional downregulation of *UNG* and *REV1* observed in Figure 3.

We further confirmed that these mutations were driven by AID rather than random polymerase errors. Mutation frequency analysis revealed a ∼3-fold increase in mutation rates targeted to canonical AID targeting motifs (WRCH hotspots) (Fig. 4C). Visualizing the mutational landscape of light chain sequences confirmed a high density of mutations concentrated within the antigen binding complementarity determining regions (CDRs) (Fig. 4D). Together, these data suggest XPO5 loss creates a permissive environment for the accumulation of unrepaired or deleterious BCR mutations.

### XPO5 knockdown decreases global miRNA expression

To determine if the observed BCR defects stem from the canonical function of XPO5, we examined the miRNA landscape. While XPO5 is the established nuclear exporter of pre-miRNAs, its absolute essentiality is a subject of debate.^41^ Studies have shown the knockout of XPO5 induces a global reduction in miRNA expression, yet enigmatically, others suggest alternative pathways of miRNA biogenesis. However, given that numerous miRNAs are critical to B cell development and function, we hypothesized XPO5 regulates B cell function in a miRNA-dependent manner.^33,48,66,67^

We performed small RNA sequencing to assess the impact of XPO5 depletion. Results indicated a global downregulation of miRNA expression at a median log2 fold change of -0.438 (Fig. 5A). Notably, we observed miRNAs that are highly expressed in control cells displayed the strongest trend towards down regulation (Fig. 5B). This suggests that nearly all miRNAs in Ramos B cells depend on XPO5-mediated export.

**Figure 5.**
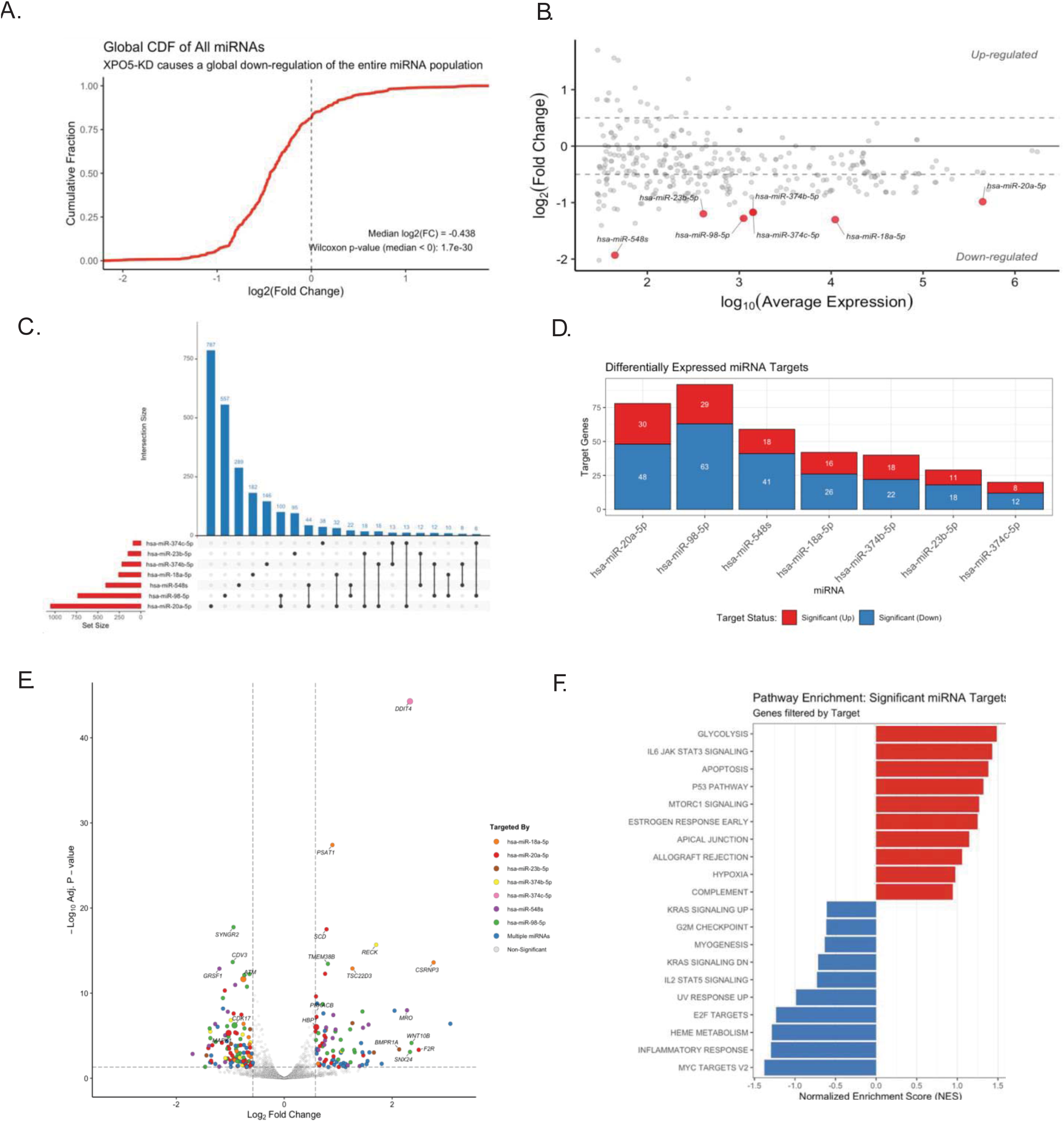
XPO5 knockdown decreases global miRNA expression. (A) Cumulative distribution function (CDF) of log2 fold changes for all detected miRNAs. Significant leftward shift indicates a global decrease of mature miRNA expression in XPO5 knockdown cells (Wilcoxon rank-sum test, p < 0.001). (B) MA Plot displaying relationship between average abundance (log10) and differential expression (log2 fold change). Top differentially expressed miRNAs are labeled. (C) Quantification of total target genes of the top 7 differentially expressed miRNA overlap between miRNA pairs. Horizontal (red) bars display total number of predicted target genes. Vertical bars (blue) represent number of gene targets for individual miRNA or overlap of miRNA pairs. (D) Differential expression of predicted target genes. Stacked bars show total number of significantly up (red) or down (blue regulated in the bulk RNA-seq data. (E) Volcano plot of differentially expressed genes in XPO5 knockdown cells. Transcripts colored according to the specific miRNA it is targeted by. (F) Gene Set Enrichment Analysis (GSEA) of differentially expressed miRNA target genes. Normalized Enrichment Scores (NES) show pathways that are activated (red) or suppressed (blue) in XPO5 knockdown cells.

Next, we aimed to understand the connection between miRNA loss and BCR expression as well as fitness defects. Here, we contextualize small RNA sequencing with our bulk RNA sequencing results. Focusing on the 7 significantly differentially expressed miRNAs, we identified a wide regulatory network comprised of >2,000 target genes (Fig. 5C). Combined, these miRNAs directly target 130 significantly upregulated genes and 230 significantly downregulated genes within the bulk RNA dataset (Fig. 5D). The significantly upregulated genes are likely a result of direct interaction where target genes are no longer repressed by these miRNAs, while the down regulated genes may reflect secondary or indirect interactions.

Next, we investigated whether the dysregulation of specific miRNA target genes could account for the functional defects observed in XPO5 depleted cells. In line with the observed G1/S arrest, we found the upregulation of miRNA target genes DDIT4^68,69^ and HBP1^70,71^, acting as mTORC1 and transcriptional repressors, respectively, prevent cell cycle at this stage. In contrast, MAPK1^72,73^ and CDK17^74,75^, required for mitogenic signals and cell cycle checkpoint progression, were downregulated (Fig. 5E). Cell fitness defects and BCR loss may also be attributed to DNA damage through an increase in AID mediated mutation. The catalytic subunit PRKACB^76^, responsible for the phosphorylation and activation of AID, is upregulated while ATM^77,78^, a key component of the DNA damage response, is down regulated suggesting an imbalance between mutagenic activity and DNA repair capacity (Fig. 5E).

Finally, we performed pathway enrichment analysis of the miRNA gene targets to identify broad trends of XPO5 regulation. This analysis identified a general shift away from proliferation and toward a stress responsive state. Consistent with the observed cell cycle defects and increased hypermutations, we observed a suppression in growth driving pathways such as MYC Targets, E2F Targets, G2M Checkpoint alongside the activation of apoptotic signaling pathways, such as P53 Pathway and Apoptosis (Fig. 5F). Overall, this suggests a multi-faceted role for XPO5 mediated miRNA regulation, driving B cell proliferation while maintaining BCR surface expression, potentially by maintaining the DNA repair threshold required to tolerate somatic hypermutations.

## Discussion

In this study, we systematically identified regulators of BCR surface expression using a genome-wide CRISPRi screen, revealing XPO5 as a factor for maintaining BCR surface expression and functional integrity. XPO5 depletion accelerated the expansion of a BCR negative population, reduced proliferation, and altered DNA repair within in the light chain. These phenotypes occur without major transcriptional changes to the BCR or accessory components suggesting a post-transcriptional mechanism of regulation.

XPO5 was originally identified as a member of the karyopherin-β family responsible for RanGTP-dependent export of double-stranded RNA-binding proteins.^79,80^ Currently, it is recognized as the canonical nuclear exporter of pre-miRNAs during miRNA biogenesis due to its structural binding preferences. Specifically, XPO5 binds to the 2-nucleotide overhang on 3’ ends of pre-miRNAs, effectively shielding them from nuclear degradation while facilitating their export for further Dicer processing.^81,82^ However, the redundancy of XPO5 has been questioned. Contrary to earlier models citing the necessity of XPO5 in miRNA biogenesis, recent studies have demonstrated that XPO5 knockout leads to a global decrease in miRNA expression rather than total ablation. ^41^ Thus, rather than acting as an absolute bottleneck, XPO5 appears to act as a dominant yet non-exclusive factor, suggesting its depletion creates a specific state of cellular dysregulation, potentially in a tissue- or cell-specific manner. This context dependence is perhaps not surprising given the pleiotropic nature that miRNAs play in diverse cell types, as they do not fit cleanly into a singular pro- or anti-tumorigenic role across the numerous contexts they modulate.

The specific accumulation of C to T transition mutations in the light chain provides mechanistic evidence that XPO5 depletion impairs the base excision repair (BER) pathway. In healthy B cells, UNG removes AID generated uracils, while XPO5 depletion leads to a downregulation of *UNG* and *REV1* expression. This suggests a DNA repair bottleneck where uracils persist and are replicated as thymines. While the apparent restriction of these mutations to the light chain is intriguing, we acknowledge that RNA sequencing relies on transcript stability and may underrepresent sequences undergoing nonsense mediated decay (NMD), particularly in the heavy chain. However, the accumulation of premature stop codons and C to T transitions within light chain WRCH motifs suggests a potential failure to tolerate or repair mutations at this locus in the absence of XPO5.

The precise mechanism by which XPO5 depletion creates this mutational imbalance requires further investigation. A primary limitation of this study is the RNA-centric focus, as future studies utilizing genomic DNA sequencing will be valuable to fully map the mutation spectrum independent of transcript stability or expression-based biases. Alternatively, this bias could be biological, potentially involving differential chromatin accessibility or chain specific recruitment factors in the absence of miRNA regulation. Additionally, while our data suggest a mutation driven loss of BCR surface expression, we did not explicitly assess protein folding or trafficking defects. Further work must also be performed to clarify the impact of individual miRNA function to better understand the regulatory role of XPO5. Finally, future in vivo studies in primary B cells will also be essential to determine the effects of XPO5 on SHM dynamics and cell homeostasis during a physiological immune response.

Our study also highlights a broad role for XPO5 in post-transcriptional regulation via miRNA control. Consistent with its established function, XPO5 depletion caused a global decrease in mature miRNA expression, highlighting its requirement for critical miRNA species. Integration of this data with bulk RNA sequencing revealed the upregulation of miRNA target genes that drive cell cycle arrest (*DDIT4*, *HBP1*) and the suppression of DNA damage response factors (*PRKACB*, *ATM*). This suggests a model where XPO5 acts to maintain high levels of miRNA expression required for proper AID activity and DNA repair (Fig. 6). Without these miRNA species, the cell enters a stressed state characterized by a repair deficit relative to mutagenic load, leading to the loss of BCR surface expression. Curiously, despite the global suppression of the miRNA landscape, our sequencing revealed a subset of miRNAs, including miR-1 and miR-2, was paradoxically upregulated. This observation may be explained by differential stability or half-life. If most of the miRNA pool turns over rapidly and cannot be replenished due to XPO5 depletion, species with a longer half-life will appear upregulated.^83^ Alternatively, these specific miRNAs may utilize non-canonical export mechanisms. Previous studies have shown miRNAs associated with stress responses or possessing distinct structural caps, such as miR-320, can utilize the Exportin-1 (XPO1) export pathway.^84,85^ In the context of cell stress induced by XPO5 knockdown, the cell may preferentially engage alternative pathways to maintain threshold levels of critical miRNAs.

**Figure 6.**
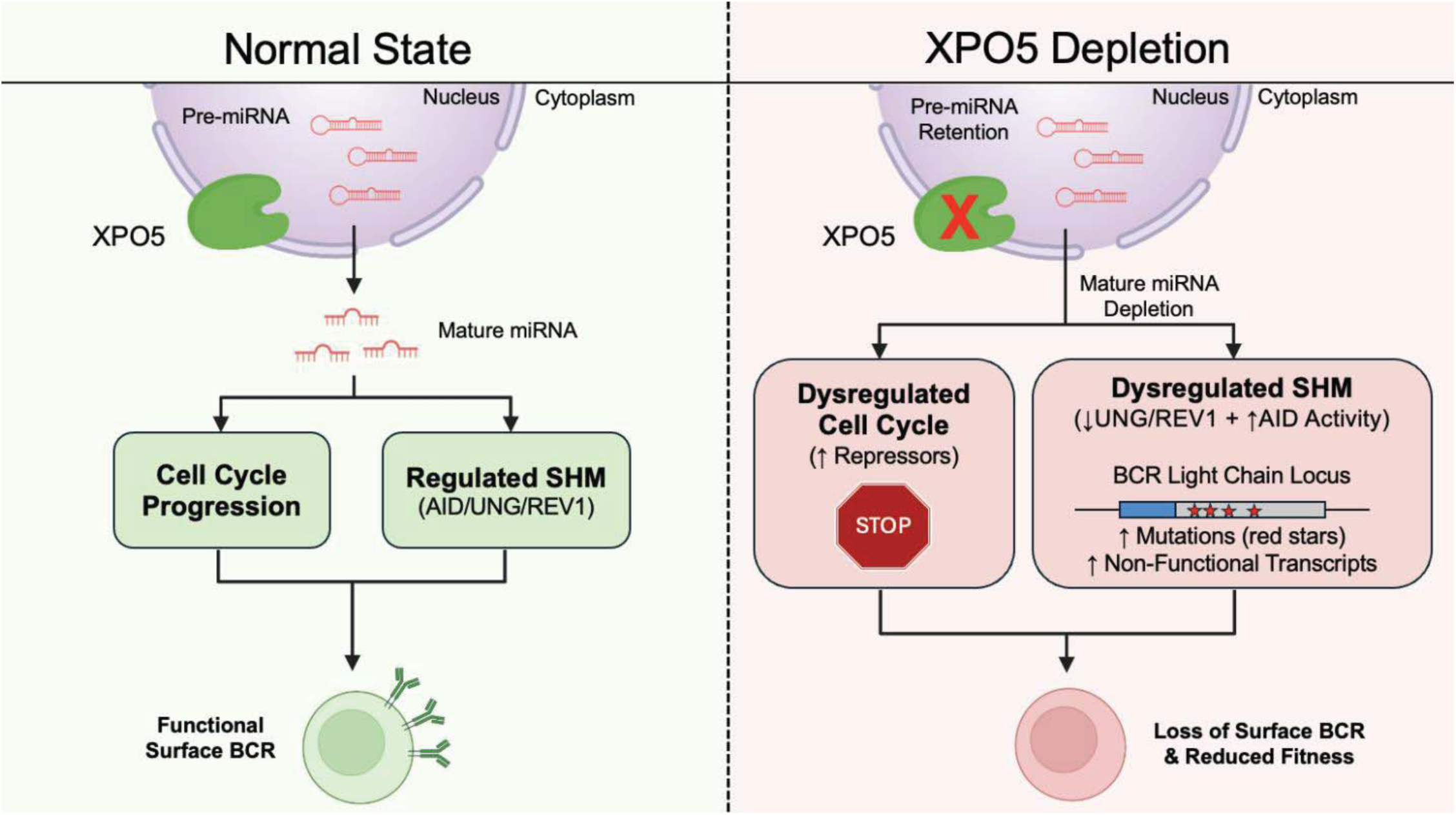
Proposed model of XPO5 mediated regulation of BCR surface expression and cell fitness. (Left) In the homeostatic state, XPO5 facilitates the nuclear export of pre-miRNAs, maintaining the pool of mature miRNAs necessary for regulating cell cycle progression and balanced SHM dynamics. This ensures the generation of functional BCR transcripts required for surface expression. (Right) Upon XPO5 depletion, pre-miRNAs are retained in the nucleus, leading to a global depletion of mature miRNA. The loss of miRNA mediated post-transcriptional regulation results in the upregulation of cell cycle repressors and dysregulated DNA repair mechanisms. These drive unbalanced SHM dynamics, specifically biasing mutations towards the BCR light chain, generating non-functional transcripts leading to the loss of BCR surface expression.

Additionally, the observed proliferation defects may be due to dual factors. First, the loss of BCR surface expression might impair tonic signaling required for peak B cell fitness. Second, the loss of direct miRNA regulation of cell cycle progression. At this time, we cannot conclude the exclusivity for either variable but suggest these may be synergistic effects stemming from XPO5 depletion. This highlights XPO5 as a crucial factor for general B cell homeostasis whose loss renders cells functionally compromised.

These findings extend beyond basic B cell biology to lymphoma pathogenesis. Dysregulation of the BCR and SHM machinery are hallmarks of B cell malignancy and autoimmunity. By identifying a link between XPO5 mediated miRNA regulation and BCR integrity, this work suggests perturbations in miRNA nuclear export machinery may be an underappreciated area of B cell biology.

## Methods

### Cell Lines and Cell Culture

The human Burkitt Lymphoma Ramos cell line was used for all tests. Ramos cells were cultures in complete RPMI-1640 and supplemented with 10% fetal calf serum (FCS), 1X penicillin-streptomycin. Cells were cultured at or below 1x10^6^ cells/mL and maintained at 37C in a humidified 5% CO2 atmosphere.

### Lentivirus Production

To generate lentivirus, HEK293T cells were seeded in DMEM with 10% FCS and penicillin-streptomycin 24 hours prior to transfections. Cells were transfected with lentiviral packaging plasmids pMD2.G (Addgene #12259) and psPAX2 (Addgene #12260) along with lentiviral transgene vector using PEI. Lentiviral supernatants were collected 72 hours post transfection and cleared via centrifugation at 800g for 10 minutes. Cleared supernatants were filtered using a 0.45um filter for immediate use or storage at -80C.

### Genome-Wide CRISPRi Screen

To generate dCas9-KRAB expressing Ramos cells, Ramos cells were transduced with dCas9-KRAB-mcherry (Addgene #188769) and expanded for 3 days. dCas9-KRAB-mCherry+ cells were sorted using the Sony Cell Sorter SH800Z and subsequently single cell cloned by limiting dilutions. A single clone stably expressing high levels of dCas9-KRAB-mCherry was expanded to obtain the CRISPRi Ramos cell pool.

300 million Ramos cells stably expressing dCas9-KRAB-mCherry were transduced with the lentiviral-based pooled hCRISPRiV2 library (Addgene pooled library #1000000090) containing 209,070 sgRNA sequences at a multiplicity of infection (MOI) of 0.1, followed by puromycin (1ug/mL) selection 48 hours post transduction. Pooled library cells were passaged every 72 hours, returning cells numbers to at least 200 million total cells to maintain sgRNA representation. After 14 days, cells were stained for surface expression of IgM using FITC-conjugated anti-human IgM antibody (Biolegend) and sorted with the Sony Cell Sorter SH800Z. In total, 200 million cells were sorted in duplicate for the highest and lowest 10% surface IgM expression. 200 million cells were harvested prior to the cell sort to act as the pre-sort control.

Genomic DNA from all samples (BCR^pos^, BCR^neg^, and pre-sort) was extracted using the Monarch Genomic DNA Purification Kit according to manufacturer’s instruction. sgRNA sequences were PCR amplified using Q5 PCR Master mix (NEB) and subsequently re-amplified with sequencing indexes. The resulting libraries were sequenced using the Illumina Hiseq platform.

### CRISPRi Screen Data Analysis

Raw sequencing reads were processed using the MAGeCK software package. First, “mageck count” was used to trim adapter sequences and align to the hCRISPRi-v2 library reference to generate a count table for all sgRNAs. To identify significant regulators of BCR surface expression, the “mageck test” function was performed to compare sgRNA abundance between BCR^neg^, BCR^pos^, and presort populations. Gene level significance was determined using the Robust Rank Aggregation (RRA) method to identify significantly enriched or depleted sgRNA sequences.

### Individual CRISPRi Knockdowns

To validate CRISPRi screen results, individual sgRNA sequences targeting desired human genes were obtained from the CRISPRiV2 sgRNA library. sgRNA oligos were obtained from Integrated DNA Technologies (IDT) and cloned into the pLentiGuide-Puro vector (Addgene plasmid #52963) via BsmBI restriction sites. Cells were transduced with individual sgRNA lentivirus and subsequently selected by puromycin (1ug/mL) 48 hours post-transduction. To ensure constant expression of the sgRNA transgene, cells were continuously passaged in puromycin.

### Flow Cytometry

200,000 cells were pelleted by centrifugation at 500g for 5 minute and resuspended in 50uL FACS buffer (1% BSA, 2 mM EDTA, PBS) and stained with the following anti-human antibodies for 30 minutes on ice protected from light: IgM-FITC, CD19-APC, B2M-APC, CD45-APC, CD20-biotin. Cells were rinsed twice by adding 500uL FACS buffer and centrifugation at 500g for 5 minutes. Pellets were resuspended in 100uL of FACS buffer for analysis. Biotin antibody-stained samples were stained again using streptavidin-APC using the same staining procedure. Data was collected using the MACSQuant 10 (Miltenyi) or sorted using the Sony Cell Sorter SH800z and analyzed using Flowjo software.

### Bulk RNA-Seq

5 million cells were pelleted by centrifugation at 500g for 5 minutes prior to RNA purification using the Zymo Quick-RNA Miniprep kit according to manufacturer’s instruction for total RNA (Zymo Research). RNA was reverse transcribed with PCR adaptors added by template switching. cDNA was amplified and prepared for sequencing using the Nextera XT kit following the manufacturer’s protocol (Illumina). Libraries were sequenced using the Hiseq Illumina next generation sequencing at a read depth of 30 million reads per sample. Sequencing data was aligned to the human reference genome (hg38) using STAR. Gene counts were generated using featureCounts and subsequent differential gene expression analysis was performed using DESeq2 in R with a significance cutoff of adjusted p < 0.05.

### Small RNA-Seq (miRNA-seq)

5 million cells were pelleted by centrifugation at 500g for 5 minutes prior to RNA purification using the Zymo Quick-RNA Miniprep kit according to manufacturer’s instruction for small RNA (Zymo Research). RNA for all samples sent to Azentia for small RNA sequencing at a read depth of 30 million reads per sample. Raw reads were adapter trimmed using Cutadapt as reads are between 18-30 nucleotides. Alignment to the human reference genome (hg38) was performed using STAR with parameters optimized for small RNA. Due to high similarity between miRNA within the same cluster, local alignment and multimapping was allowed, while splicing was disabled. Mature miRNA counts were quantified using featureCounts guided by miRbase v22 annotation. Differential expression was determined using DESeq2 in R.

### BCR Sequence Reconstruction and Mutation Analysis

To identify somatic hypermutation (SHM) variants, the dominant Ramos BCR heavy and light chain variable region sequences were reconstructed from bulk RNA-seq data using TRUST4. The consensus V(D)J and VJ sequences were added to the standard human reference genome (GRCh38) to create a custom reference. RNA-seq reads were re-aligned to this custom reference using STAR.

Mutation calling was performed using a custom script utilizing SAMtools. Pileup files were generated using “samtools mpileup” restricted to the heavy and light chain sequences, with a minimum mapping quality (MAPQ) of 30 and minimum base quality 30. A custom python script was used to parse pileup data and calculate mutation frequencies. Variants were filtered using a consensus approach, requiring support from both forward and reverse strands with a minimum of 3 total reads.

### Cell Proliferation and Competition Assay

To generate GFP control cells, dCas9-KRAB-mCherry non-target sgRNA expressing Ramos cells were transduced with constitutive GFP expression lentivirus. 3 days post transduction, cells were sorted for GFP expression using the Sony Cell Sorter SH800z. Cells were seeded at a 1:1 ratio of GFP non-target cells to non-target or XPO5 targeting sgRNA expressing cells for a total of 200,000 cells (100,000 cells per cell type) in 6-well plates to allow sufficient room for expansion. Cells were observed daily starting on the day of initial plating through 10 days for GFP percentages using the MACSQuant 10 (Miltenyi). GFP negative and GFP positive cells were used as controls to distinguish the two populations.

### Cell Cycle Analysis

Prior to cell cycle analysis dCas9-KRAB-mCherry Ramos cells expressing desired sgRNA knockdown sequences were plated in triplicate at 100,000 cells/mL in 6-well plates for a total of 5mL suspension. After 24 hours, cells were harvested and rinsed using PBS prior to staining with Hoechst 33342 dye diluted to 10ug/mL in PBS. Cells were incubated protected from light at 37C, 5% CO2 for 15 minutes. Excluding doublets, percentages of single cells in G1, S, and G2-M phases were determined using the MACSQuant 10 (Miltenyi). Unstained control was performed in cells without Hoechst staining.

### Anti-IgM Apoptosis (Cell Death) Assay

dCas9-KRAB-mCherry Ramos cells expressing desired sgRNA knockdown sequences were plated in triplicate at 100,000 cells/mL in a 24-well plate prior to stimulation with anti-IgM at concentrations of 0ug/mL, 3 ug/mL, and 5ug/mL. After 24 hours, cells were washed once with PBS buffer and then stained with 300nM DAPI solution for 5 minutes. Cells were washed twice with PBS and then analyzed using the MACSQuant 10 (Miltenyi).

### Statistical Analysis

Data are presented as mean +/- standard deviation (S.D.) of at least n=2 biological replicates, as indicated in figure legends. Statistical significance for pairwise comparisons was determined by an unpaired, two-tailed Student’s t-test. For global miRNA distribution, a Wilcoxon rank-sum test was performed. Gene Set Enrichment Analysis (GSEA) was performed using rank gene lists. P < 0.05 were considered statistically significant.

## Acknowledgments

We thank David Krag and Stephanie Pero for providing access to essential equipment and technical resources that made this work possible. We also thank Jonathon Boyson for his valuable guidance and support throughout the research process.

## Data Sharing Plans

All data generated in this study will be deposited in the NCBI Gene Expression Omnibus (GEO) and will be accessible to the public as of the date of publication. All custom code and scripts used for data analysis will be made available via GitHub repository upon publication.

## Author Contributions

T.K., H.P., and D.M. designed the research. T.K. performed research. T.K., Z.M., J.D., W.D., S.L., J.C., and D.M. analyzed data. H.P. and S.F. contributed new reagents/analytic tools. All authors wrote the paper.

